# Remote manipulation of magnetic nanoparticles using magnetic field gradient to promote cancer cell death

**DOI:** 10.1101/317115

**Authors:** Mahendran Subramanian, Arkadiusz Miaskowski, Stuart Iain Jenkins, Jenson Lim, Jon Dobson

**Author notes:** Corresponding author: M. Subramanian.

## Abstract

The manipulation of magnetic nanoparticles (MNPs) using an external magnetic field, has been demonstrated to be useful in various biomedical applications. Some techniques have evolved utilizing this non-invasive external stimulus but the scientific community widely adopts few, and there is an excellent potential for more novel methods. The primary focus of this study is on understanding the manipulation of MNPs by a time-varying static magnetic field and how this can be used, at different frequencies and displacement, to manipulate cellular function. Here we explore, using numerical modeling, the physical mechanism which underlies this kind of manipulation, and we discuss potential improvements which would enhance such manipulation with its use in biomedical applications, i.e., increasing the MNP response by improving the field parameters. From our observations and other related studies, we infer that such manipulation depends mostly on the magnetic field gradient, the magnetic susceptibility and size of the MNPs, the magnet array oscillating frequency, the viscosity of the medium surrounding MNPs, and the distance between the magnetic field source and the MNPs. Additionally, we demonstrate cytotoxicity in neuroblastoma (SH-SY5Y) and hepatocellular carcinoma (HepG2) cells in vitro. This was induced by incubation with MNPs, followed by exposure to a magnetic field gradient, physically oscillating at various frequencies and displacement amplitudes. Even though this technique reliably produces MNP endocytosis and/or cytotoxicity, a better biophysical understanding is required to develop the mechanism used for this precision manipulation of MNPs, in vitro.

## I. INTRODUCTION

The manipulation of magnetic nanoparticles (MNPs) using external magnetic fields has opened up the possibility of various biomedical applications. Promising in vitro technologies have emerged, using this non-invasive external stimulus, such as gradient magnetic field assisted bio separation [Bruce 2005] and gene transfection [Scherer 2002, Subramanian 2013]; alternating field gradient mediated cytotoxicity [Hapuarachchige 2016]; dynamic magnetic field (DMF) mediated cytotoxicity [Zhang 2014]; alternating magnetic field (AMF) mediated cytotoxicity [Subramanian 2016]; and controlled drug release [Satarkar 2008]. The techniques mentioned above involve both permanent magnets and electromagnets with different working principles. Bioseparation consists of the use of a field gradient to capture specific biomolecules which are bound to MNPs [Dutz 2016]. Magnetofection involves the use of a magnetic field to attract magnetic nanoparticles and nucleic acid complexes towards cells to facilitate gene transfection [Mykhaylyk 2007, Subramanian 2017]. The alternating field gradient mediated cytotoxicity technique uses an alternating field gradient (Gz- 95 G/cm) within a homogenous field (a 9.4 T preclinical MRI system) to align the nanoparticles parallel to the homogenous field and destroy cancer cells through an induced motion of magnetic nanoparticle aggregates within cells [Hapuarachchige 2016]. Dynamic magnetic fields (10 to 20 Hz, 30 mT) encourage torques, i.e., repeated incomplete rotational movements, of MNPs which enables remote the stimulation of cell death by the permeabilizing of the lysosome membrane and the triggering of apoptosis (programmed cell death) [Zhang 2014]. The application of alternating magnetic fields (50 - 1000 kHz) to MNPs which are bound to receptors on cell membranes or internalised within cells, has been used to activate chemical signalling in the cell, leading to apoptosis or the depositing of energy, so triggering necrosis (cell death or tissue damage in an organ) [Subramanian 2016].

The possible physical explanations for the static magnetic field gradient mediated attracting/aligning of MNPs towards a magnet field have been discussed extensively [Pankhurst 2003, Furlani 2008, Furlani 2012, Garraud 2016]. Moreover, a possible mechanism for alternating field gradient mediated cytotoxicity has been examined [Hapuarachchige 2016]. However, an explanation for dynamic field induced rotational motion has proven elusive; it is stated in [Zhang 2014] that a dynamic field generates a torque, equal to and that this enables the rotation of individual MNPs around their axis, but the dynamics of the magnetic fields used requires explanation. Finally, a deposition of energy is possible when MNPs are exposed to alternating magnetic fields in the radio frequency range since they then dissipate heat due to susceptibility, hysteresis and friction losses [Pankhurst 2003].

This study focuses on the underlying conditions behind the manipulation of MNPs using unidirectional time-varying magnetic fields/field gradients and how this can be used to induce magneto-mechanical cell death in cancer cells. The numerical model discussed here should allow researchers to further this novel technique and increase the manipulability of MNPs. Here, we have used a time varying (1 to 4 Hz) alternate pole magnet array plate populated with Nd-Fe-B permanent magnets (~ 450 mT) to create field gradients which result in the enhanced attraction of magnetic nanoparticle towards cells and so the induction of cytotoxicity in neuroblastoma (SH-SY5Y) and hepatocellular carcinoma (Hep G2) cancer cells.

## II. MATERIALS AND METHODS

### A. Magnetic field measurements

Magnetic field measurements of three different 3 × 3 Nd-Fe-B magnet arrays were performed using a Hall probe (F.W.Bell - 5080 Teslameter; Orlando, Florida, USA). The numerical modeling discussed in this paper was carried out using the Sim4Life (ZMT Zurich MedTech, Zurich, Switzerland) platform. The dimensions of the magnets and the corresponding distances were measured in relation to the magnet array which was explicitly fabricated for performing the experiments here.

### B. Mathematical model

These measurements were used to calculate the magnetic field (flux density, T) of each 3 × 3 array (Fig. 1), the magnetic force (N) experienced by the MNPs, and the gradient field (gradient flux density; T/m) for specific distances from the magnet faces along the z-axis (0 mm, 1 mm, 1.5 mm) and from the centre of each magnet along the y-axis (−3 to +3 mm; 2D plot).

**Fig. 1.**
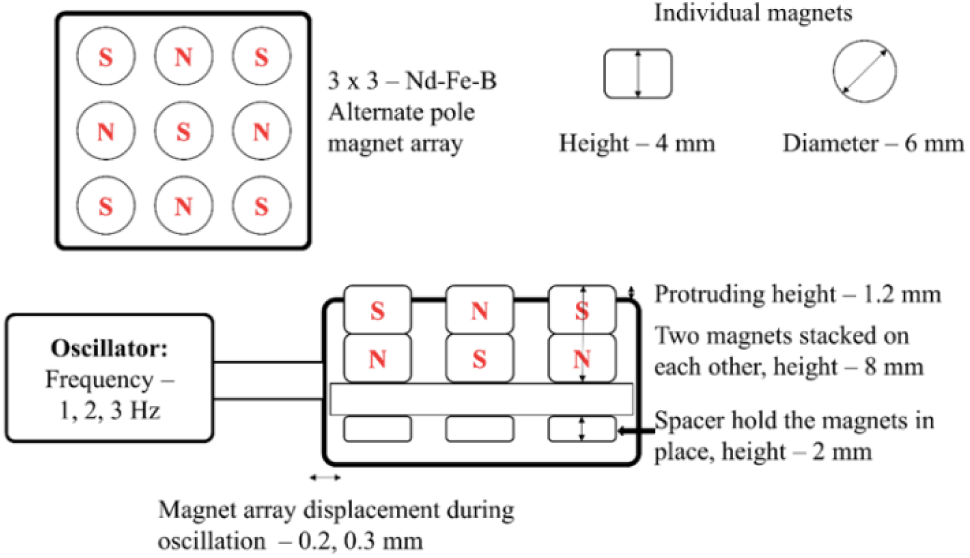
Illustration of the alternate pole magnet array, made up of Nd-Fe-B permanent rare earth magnets, which was used for the induction of magneto mechanical cytotoxicity in cancer cells. Dimensions of the magnet used (height, diameter) and the populated magnet array (cross section) connected to the oscillator are provided.

The magnetic field was calculated using the magnetic vector potential (***A***) under the assumption that ∇ · (***B***) = 0 and *B* =∇ × (4), i.e.:

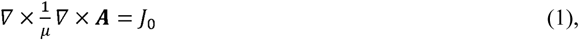
where *J*_0_ is the surface current density (A/m).

In addition, in order to explain the aforementioned phenomenon (magneto-mechanical cell death), the following mathematical model can be used.

The magnetic moment, **m**, of a nanoparticle (based on the typical design of having magnetic material at the core) is a product of its magnetization, **M**, and volume, 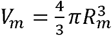, where *R_m_* is the radius of its magnetic core:

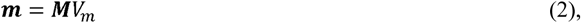

The volumetric magnetization of the MNPs is induced by application of the external magnetic field, ***H***,

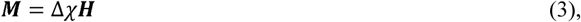
where ∆*χ* is the effective susceptibility of the MNPs with respect to the medium (water) and

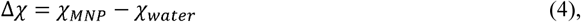

Assuming that the relative permeability of the water, *μ_r_* = 1, together with ***B*** = *μ*_0_***H*** the volumetric magnetization can be introduced as

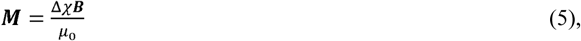
and

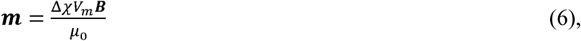

Furthermore, if MNPs are point dipoles with the same moment, **m**, the force experienced by them is

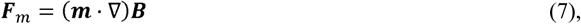

Combining (6) and (7) we get

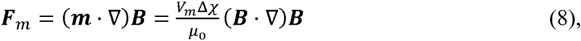

Knowing that ∇ × ***B*** *=* 0, the force on the dipole can be modified as

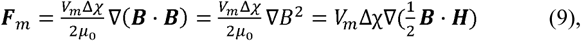

To be effective, **F**_m_ must overcome the hydrodynamic drag force, **F**_d_, acting on the MNPs, i.e.

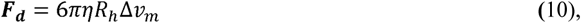
where *η* is the viscosity of the medium (water) surrounding the MNPs, *R_h_* is the hydrodynamic radius, and *v_m_* is the drift velocity (∆*v_m_ = v_MNP_ − v_water_*). The MNPs will be immobilized when **F**_d_ = **F**m, so

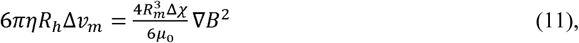
or

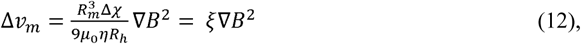
where ξ is the magnetophoretic mobility.

As the first approximation, it can be assumed that *R_m_ = R_h_* and when

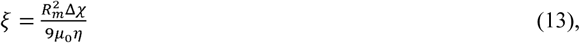

As the second approximation, it can be assumed that *R_m_* ≠ *R_h_* and when

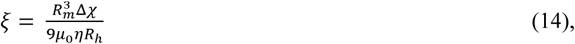
where 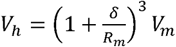 and *δ* is the thickness of the absorbed surface layer.

### C. Cell culture

Human neuroblastoma cells (SH-SY5Y; CRL-2266; American Type Culture Collection, Manassas, USA) were maintained as described previously [14]. SH-SY5Y cells were seeded at a density of 1 or 2 × 10^4^ cells in 100 μl medium onto uncoated 96-well plates in Ham’s F12: MEM (1:1) supplemented with 10% fetal bovine serum (FBS), 2 mM L-glutamine, 100 U penicillin, and 0.1 mg/ml streptomycin. Human hepatocellular carcinoma cells (Hep G2; HB-8065; American Type Culture Collection, Manassas, USA) were cultured in DMEM with 10% FBS, 2 mM L-glutamine, 100 U penicillin, and 0.1 mg/ml streptomycin, prior to seeding at a density of 1 or 2 × 10^4^ cells in 100 μl medium onto rat tail collagen I-coated 96-well plates. Cultured cells were incubated at 37 °C, 5 % CO_2_, for 24 h prior to cytotoxicity experiments.

### D. Magnetic field gradient mediated cytotoxicity experiments

Magnetite nanoparticles were obtained from Ozbiosciences, Marseilles, France (Polymag MNPs, used with SH-SY5Y cells) and nanoTherics Ltd, Staffordshire, UK (nTmag MNPs, used with Hep G2 cells). Both Polymag and nTmag contains a magnetite core. They range between 100–250 nm in hydrodynamic diameter and are surface functionalized with proprietary polyethyleneimine derivative. Both have positive zeta potential (surface charge). The rationale for using both particles is that the study was aimed at evaluation of MNP + magnet array mediated cancer cytotoxicity. Our goal was to demonstrate that the technology works efficiently with different commercially available magnetic nanoparticles that have been widely demonstrated to show high biocompatibility [Subramanian, M., et al., 2017; Fouriki, A. and Dobson, J., 2014]. MNPs were prediluted in 100 μl medium and added to cell cultures (0.2 μl per well) immediately prior to magnetic field application. Cell culture plates form fit on top of the magnet array (cells were above magnet surface, 1 mm along z axis) and the magnet array was moved back and forth along the x axis using a stepper motor for 30 minutes with the indicated frequencies and displacements. The experiment was conducted within an incubation hood during the 30-minute exposure.

### E. Cell viability assay

After exposure to the oscillating magnet array, cell culture plates were returned to the incubator for 48 hours before viability testing. The CytoTox-ONE^TM^ membrane integrity assay (Promega, Southampton, UK) provided a measurement of the amount of lactate dehydrogenase (LDH) released through damage to cell membranes. The assay was performed according to the manufacturer’s instructions, and luminescence was recorded using a plate reader (BioTek, Bedfordshire, UK).

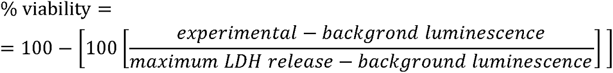

### F. Magnetic nanoparticle uptake

0.2 μl Polymag nanoparticles were incubated with 0.2 μg pEGFPN1 plasmid (encoding the green fluorescent protein, GFP) for 15 mins, then added to cultures which were immediately placed on the magnet array (30 minutes oscillation at 2 Hz frequency, 0.2 mm horizontal displacement). GFP expression indicates cellular uptake of nanoparticles and expression of the plasmid. Fluorescence was observed after 48 hrs incubation using a Leica fluorescent microscope – DM series (Newcastle Upon Tyne, UK).

### G. Statistics

If not stated otherwise, the data were presented as mean ± SD of values from at least three experiments. Statistical significance was determined by a one-way ANOVA followed by a Bonferroni post hoc test (multiple comparison test) with *p* > 0.05 considered as not significant.

## III. RESULTS

The magnetic field surrounding an array of magnets was measured to inform a theoretical prediction of magnetic nanoparticle behavior within such a field. Cancer cell lines were exposed to magnetic nanoparticles in the presence of an oscillating magnetic field gradient to assess their potential for inducing cytotoxicity.

### A. Magnetic field measurements

Magnetic field (flux density) measurements above the magnet array were determined at the magnet face, and base of a culture plate immediately above the array (see Fig. 2).

**Fig. 2.**
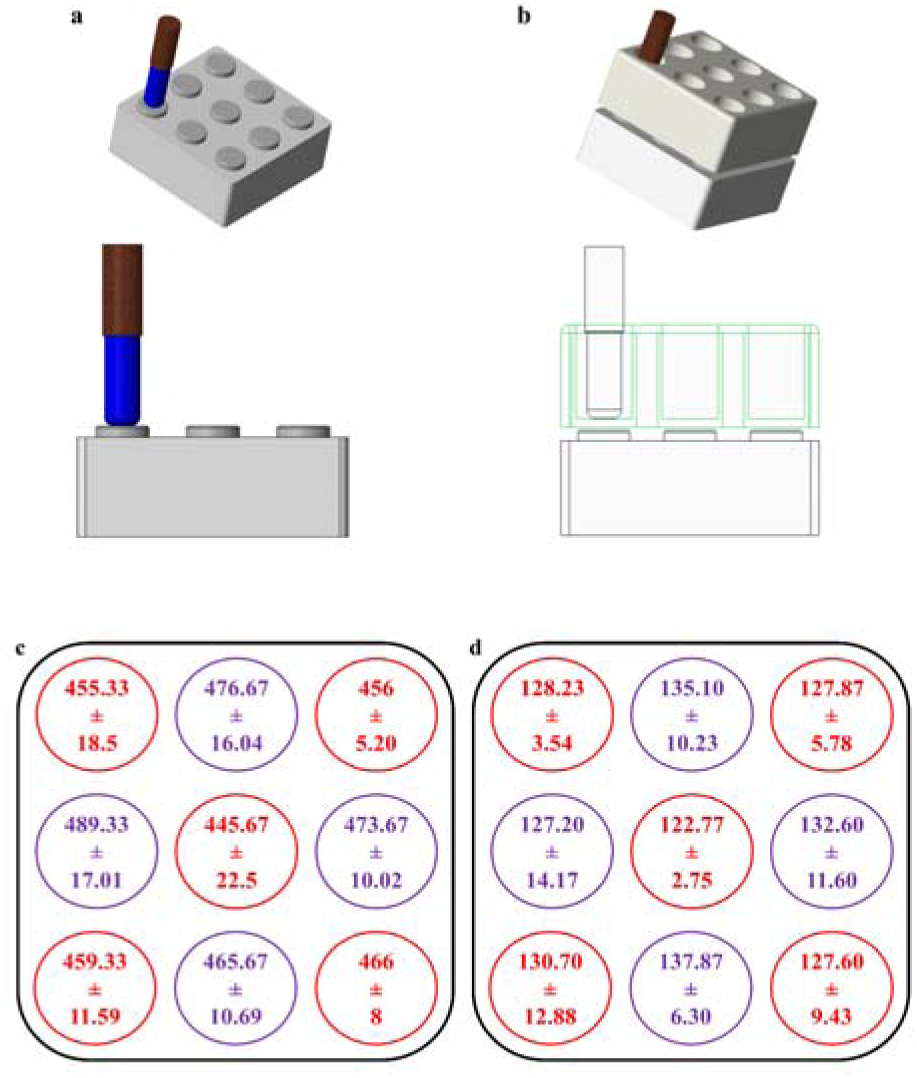
Hall probe measurements of flux density. A Hall probe (indicated as blue) was used to determine magnetic field B (flux density) at (a) 0 mm above the 3 × 3 magnet array and (b) the base of a culture plate well (1 mm above magnet face). (c) Measured field B values for method described in (a). (d) Measured field B values for method described in (b). Values are in milliTesla (mT), average of 3 measurements from 3 different magnet arrays. Red - South; Purple – North.

**Fig. 3.**
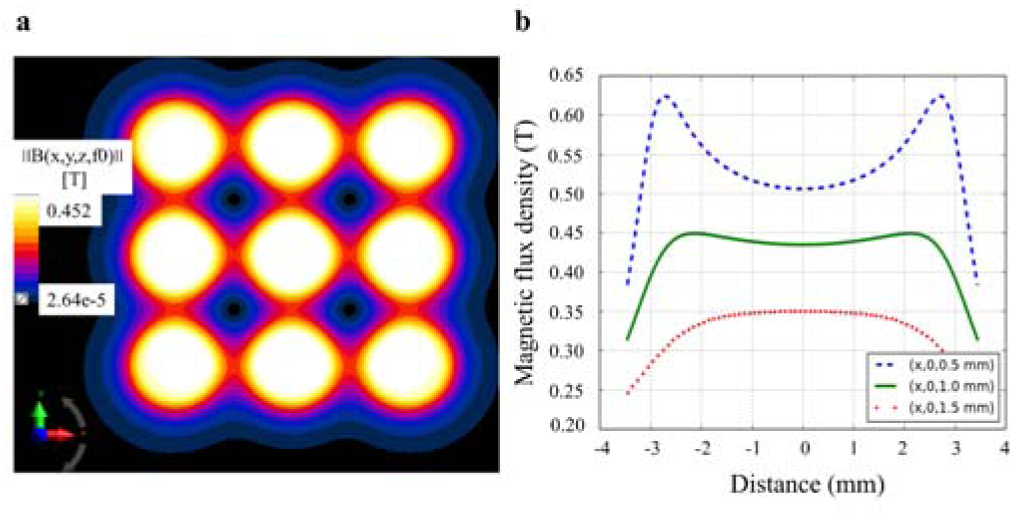
2D model of flux density above magnetic array. A magneto-static (vector potential) algorithm was used to calculate: (a) magnetic field (flux density) distribution over the 3 × 3 magnet array at 1 mm above the magnets; and (b) magnetic field (flux density) distribution at 0.5, 1, 1.5 mm above the magnets (z axis). 3 and −3 mm distance represent magnet edges. Legend shows (x, y, z coordinates in mm).

Field gradient decreases with distance (z-axis) from the magnet face (Fig. 4). Here, it is important to note that the internal diameter of a cell culture well is 6.5 mm (typical 96 well plate). Figure 4(b) and 2d plot in 4(c) demonstrate that the magnetic flux gradient () is relatively low in the center and higher near the magnet’s edge.

**Fig. 4.**
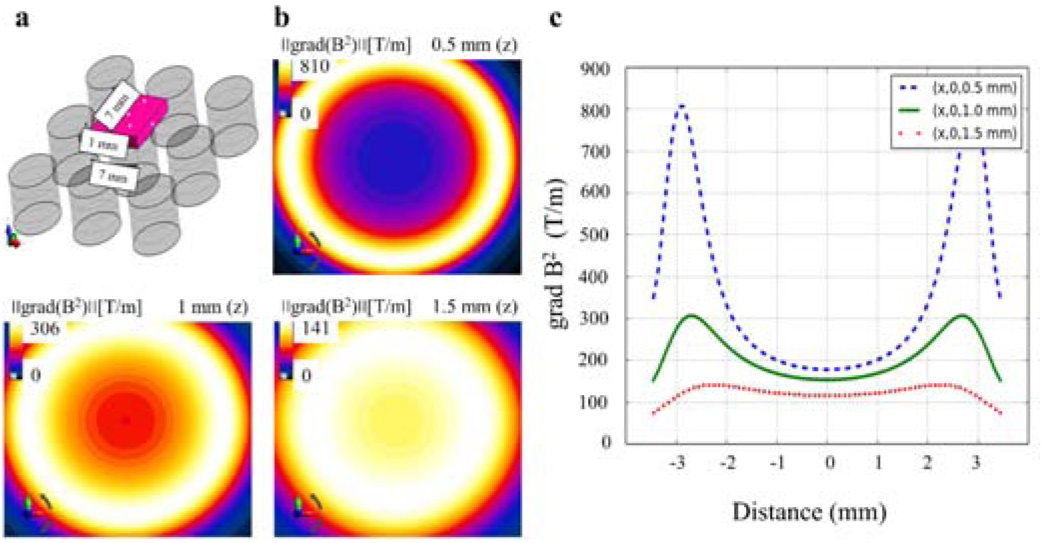
Modelling of gradient flux density above a static magnet. (a) 3D model of the 3 × 3 magnet array used for gradient flux density calculations. (b) The field gradient distribution (gradient flux density) at 0.5, 1, and 1.5 mm along the z-axis (vertically above centre of magnet, x = y = 0). (c) 2D plot of the field gradient distribution (gradient flux density) across the midline of a magnet at 0.5, 1 and 1.5 mm above the magnet face.

### B. Numerical modelling

Based on the magnetic field measurements shown in Fig. 2, magnetic field distribution and gradient flux density calculations were performed for the magnet array set up used for conducting experiments. These estimates provide insight into the principles behind the manipulation of MNPs.

The magnetic fields and the field gradients produced by the magnet array were calculated (x-, y-, z-axes). The magnetic force experienced by the MNPs and the velocity of the MNPs relative to the carrier fluid was calculated using the numerical model described in the methods section. Contour plots of the parameters mentioned above for 1 mm above the magnet array (z-axis) provide a better understanding, aiding in the positioning of the cells and MNPs (see Figs. 3, 4).

Field gradient graphs showing x-axis against y-axis (horizontal magnet displacement during array oscillation: 0 mm and 0.1 mm) should provide us with insights into the changes occurring in the magnetic field and field gradient when the magnet array is moved back and forth along the x axis (see Fig. 5). The magnetic field flux (Fig. 5a) and field gradient data (Fig. 5b) demonstrate that the magnetic field shifts in the y-axis over 0 - 0.1 mm, a relatively short distance.

**Fig. 5.**
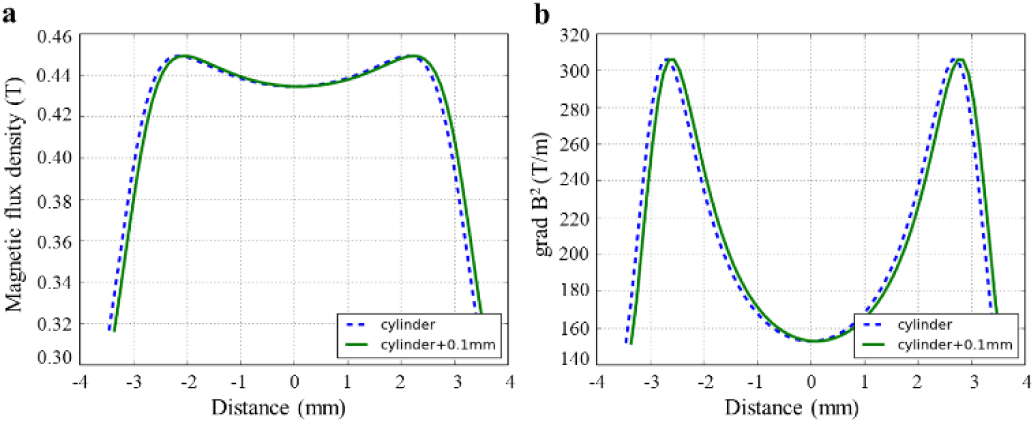
Gradient flux density at culture well surface (adherent cell layer) during horizontal oscillation of the magnetic array. (a) 2D plot of the magnetic field (flux density) for 0 mm (x axis) and 0.1 mm shifted along the y-axis. (b) The field gradient (gradient flux density) for 0 mm (x axis) and 0.1 mm shifted along the y-axis.

The magnetic force, (***F****_m_*), and the velocity of the MNPs will be directly proportional to the field gradient (**B**^2^): i.e., they will appear as a crater in 3D at 1 mm (z-axis). And the displacement of the MNPs will be inversely proportional to the time-varying frequency.

### C. Cytotoxicity of neuroblastoma cancer cells

Cytotoxicity was successfully induced in both cancer cell lines following exposure to magnetic nanoparticles and an oscillating magnetic array. The extent of cytotoxicity varied with frequency and displacement. Cell viability for SH-SY5Y neuroblastoma cells was determined through membrane integrity assay (see Fig. 6.). Viability decreased with increasing the oscillating frequency and increasing amplitude of array oscillation. Polymag MNPs, with 3 Hz/0.3 mm oscillation reduced viability to 70.5 ± 6.0 %.

**Fig. 6.**
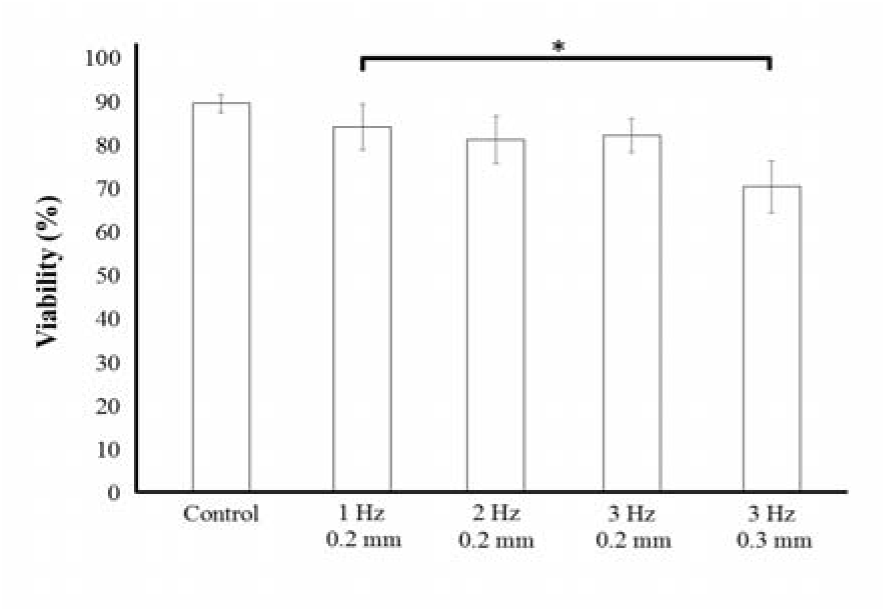
Magneto-mechanically induced cytotoxicity in SHSY-5Y neuroblastoma cells, exposed to magnetic nanoparticles and a time varying magnetic field gradient. n = 3. Control - MNPs only and no magnetic field.

### D. Nanoparticles were endocytosed by neuroblastoma cells

To establish that the nanoparticles were endocytosed by neuroblastoma cells, Polymag nanoparticles were functionalized with plasmids encoding GFP, then these complexes were incubated with the cells, in the presence of a magnetic array. Controls were incubated with plasmid only (no MNPs) but still exposed to the magnetic field. GFP expression by cells indicated endocytosis of MNPs and subsequent plasmid expression (Fig. 7a). GFP expression was never observed in control cultures. Whereas, there was no/low uptake in the plasmid only control. This demonstrates that the cytotoxicity observed (Fig. 6) can be attributed to internalization of MNPs.

**Fig. 7.**
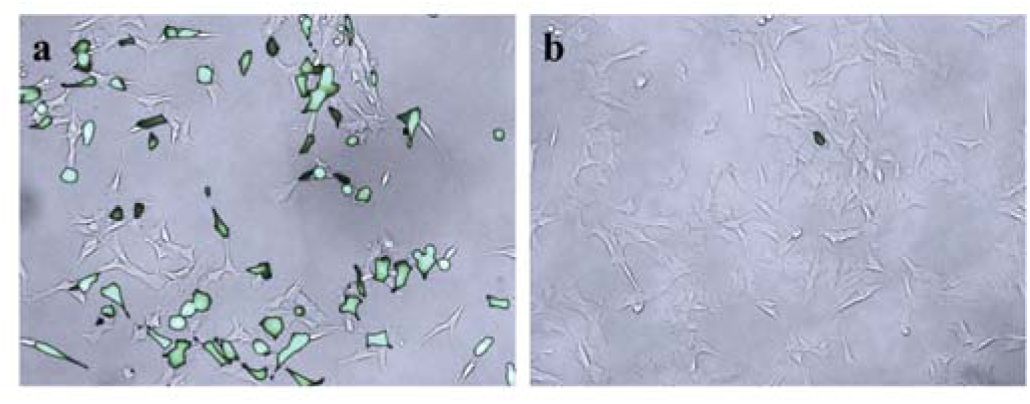
Endocytosis of Polymag MNPs by SHSY-5Y cells was confirmed by GFP expression. (a) Fluorescence micrograph of GFP+ SHSY-5Y cells superimposed on counterpart phase contrast micrograph, demonstrating expression of a GFP-encoding plasmid carried by Polymag MNPs (with oscillating magnetic field, frequency = 2 Hz, amplitude = 0.2 mm). (b) Fluorescence micrograph of GFP-SHSY-5Y cells superimposed on counterpart phase contrast micrograph, demonstrating absence of expression of GFP when incubated with plasmid without Polymag MNPs as a vector (with oscillating magnetic field, frequency = 2 Hz, amplitude = 0.2 mm).

### E. Cytotoxicity of hepatocellular carcinoma cells

At the highest oscillation frequency tested, viability of Hep G2 cells was reduced to 53.1 ± 11.4 % (Fig. 8). A decrease in viability with increasing oscillation frequency was observed 4 Hz showed significantly reduced viability versus 2 Hz.

**Fig. 8.**
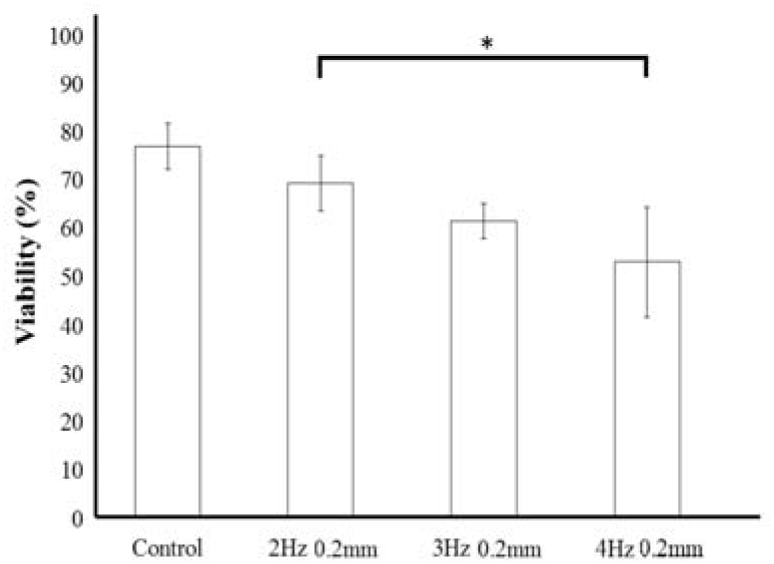
Magneto-mechanically induced cytotoxicity in Hep G2 carcinoma cells, exposed to magnetic nanoparticles and a time varying magnetic field gradient. n = 3. Control - MNPs only and no magnetic field.

### F. Magnet array optimization

Based on the observed results, we propose that magnet array design could be improved to enhance performance when delivering MNPs to a cell monolayer within a round culture well. To address this goal, we evaluated different magnet shapes and arrangements using our model, with the aim of identifying improved field gradient experienced by MNPs/cells. Two different types of magnet array were compared with the magnet array used in this study via numerical modelling. A magnet array made up of pyramidal magnets, and a custom shaped Halbach magnet array based on [Barnsley 2016, Barnsley 2017] were taken into consideration (Fig. 9). The magneto-static algorithm described earlier was used to calculate magnetic field distributions for the magnet arrays shown in figure 10. The field strength values at 1 mm above the magnet apex (z-axis) were used to calculate the field gradient.

**Fig. 9.**
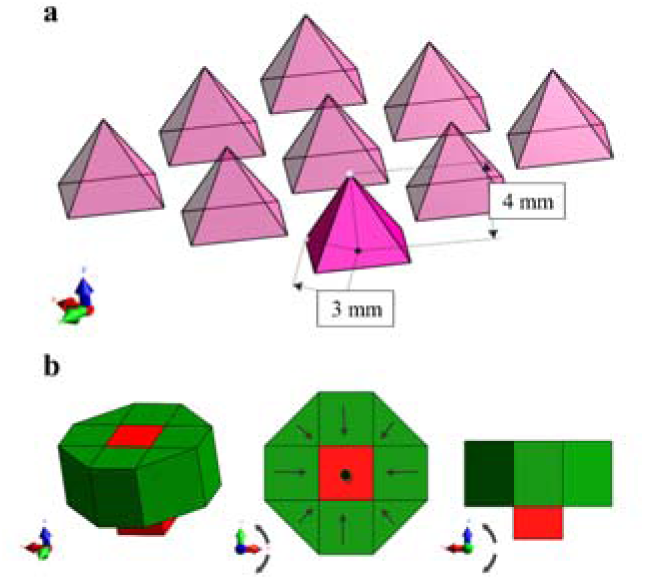
Proposed magnet array with pyramid shaped magnets (a); optimized Halbach magnet array (b).

**Fig. 10.**
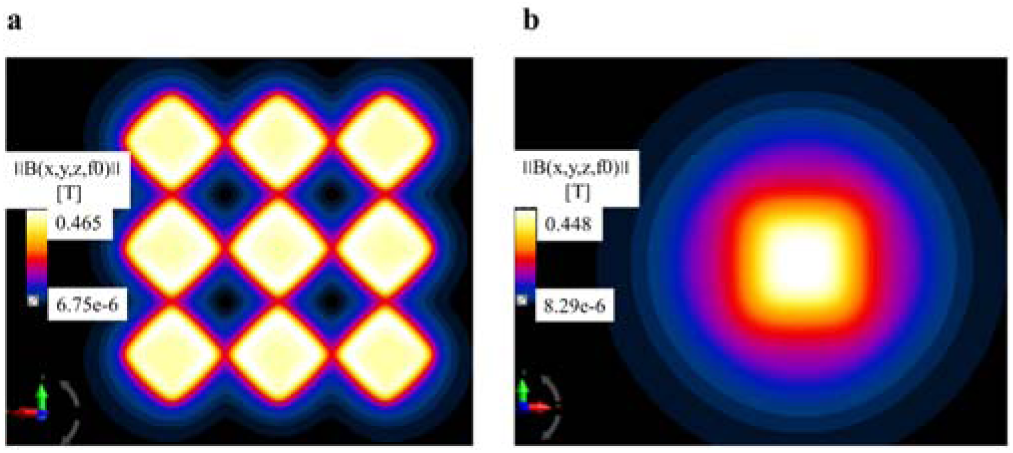
Magneto-static (vector potential) algorithm based numerical calculations: (a) magnetic field (flux density) distribution over the 3 × 3 magnet array (pyramid shaped magnets) at 1 mm above the magnets (b) magnetic field (flux density) distribution over the optimised Halbach magnet array at 1 mm above the magnets.

From Fig. 11, we can infer that the gradient field values at 1 mm and 1.5 mm (z-axis) were higher for the magnet array made up of pyramid-shaped magnets and optimized Halbach magnet array than the magnet array used in this study. Optimized Halbach array produced better gradient field out of all three designs. Moreover, is directly proportional to the gradient magnetic field (see Eq. 9). These numerical calculations are shown in figure 11 demonstrate that prospects are available for further magnet array optimization to enhance the manipulation of magnetic nanoparticles.

**Fig. 11.**
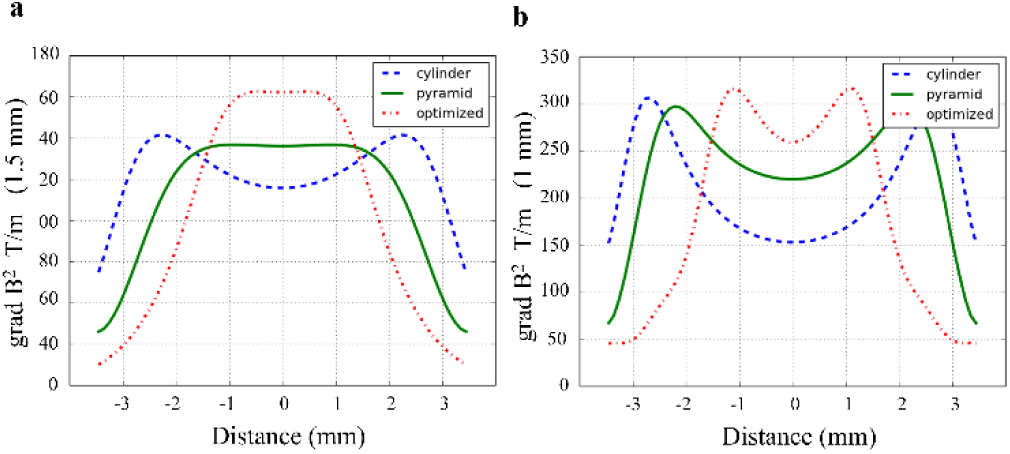
Field gradients are predicted to differ between magnets of different shape. (a) 2D plot of the magnetic field gradient (gradient flux density) at 1.5mm along the z-axis. (b) 2D plot of the magnetic field gradient (gradient flux density) at 1 mm along the z-axis.

## IV. DISCUSSION

To understand the mechanism behind the observed cytotoxicity mediated by MNPs, it is essential to study the magnetic field and the field gradients produced by the magnet array. Previous studies on a static magnetic array suggest that the magnet does accelerate the sedimentation of MNPs onto adherent cells by overcoming diffusion-limited accumulation [Furlani 2008]. This sedimentation rate can be influenced by varying the size of the MNPs and/or the distance between the magnet array and the cell culture plate [Furlani 2008]. Furthermore, a stronger axial magnetic force, achieved using an alternating magnetic pole array rather than a unidirectional magnet array, improves the accumulation rate [Furlani 2012]. There are some interesting questions posed by the results of the present study regarding the mechanism behind this unique biophysical technique which facilitates the damage of cells via a laterally-oscillating high gradient magnet array. Based on our model, the field gradient should be considered the most dynamic variable when calculating the ***F****_m_* (N) from (9). Thus, high field gradient will increase the force acting upon the magnetic nanoparticles. Assuming that the magnetic array will move without any drag or delay in the field over a displacement distance of 0.2 mm.

A scrutiny of the magnetic flux density (*B*), its squared gradient (*B*^2^), the magnetic force (***F****_m_*) and the velocity of the MNPs (Figs 2+5 and numerical calculations) suggests that these values vary within a single well (internal diameter - 6.5 mm) of the (static) cell culture vessel when an alternating-pole magnet array is moved back and forth underneath. It is already evident that the ***F****_m_* overcomes some of the viscous drag force of culture medium since the application of this alternating-pole magnet array resulted in the accumulation of 90% of MNPs at the bottom of the plate after 20 minutes of exposure to a static array [Furlani 2012]. When most of the particles are in contact with the monolayer of cultured cells, the nanoparticles will experience a field gradient and a homogenous field (Figs 3-5). MNPs in the gradient field will experience a translational motion and in the homogeneous field will experience a rotational motion - in addition to the random Brownian movement (walk) [Pankhurst 2003]. The magnetophoretic parameter (*ξ*), i.e., the manipulability of 100 nm (HD) sized MNPs used in this study, can be calculated using (14), and the displacement values resulting from the velocity do predict that the nanoparticles undergo a translational movement.

Hence, we can advocate from our calculations that the MNPs will experience oscillating horizontal translational motion when the magnet array is moved laterally (−0.1 mm to 0.1mm in the x-axis for 0.2 mm displacement or -0.15 mm to 0.15mm for 0.3 mm displacement) underneath the cell culture plate. And this displacement of MNPs is inversely proportional to the time-varying frequency of the magnet array movement. In other words, the translational movement of the MNPs, induced by the time-varying magnet array will reduce, if we increase the time-varying frequency. But, the number of movement cycles will increase.

An important point to note is that MNPs tend to form aggregates, and this may have an impact on the translational motion of the MNPs [Hapuarachchige 2016]. The low displacement values may be due to viscous damping and displacement might be higher when the nanoparticles form aggregates. However, this back and forth motion, through the influence of a time-varying field gradient, should have induced mechanical stress on the surface/inside of the cells, which in turn should increase cell death; this effect is facilitated by the magnet’s displacement. Even though the displacement values are low, the mechanical manipulation would cause stress for 30-minute exposure at 2 Hz, as the total number of oscillations would be 3600, i.e., 120 cycles per minute. And, we should not exclude nanoparticle clusters. The sizes of animal cells are usually between 0.01 mm and 0.1 mm, and cell cytoplasm has a higher viscosity than the surrounding cell culture medium. So, gradient magnetic field assisted movement of MNPs should reduce once the MNPs are endocytosed by cells; however, there are studies validating the motility of small molecules within cytoplasm [Kalwarczyk 2011].

This spatio-temporal behavior parameter, i.e., the oscillating motion of the MNPs, must be responsible for the increase in the cytotoxicity of the neuroblastoma cancer cells. *In vitro* numerical investigation suggests that magnet geometry influences the collection time of the magnetic particles moving through a high viscosity fluid. Furthermore, we use cylindrical shape magnets with an aspect ratio of 1:1.66; this is comparable to the magnet geometry used for *in vivo* studies; this is cylindrical - not conical - and has an aspect ratio of 1:1 [Garraud 2016]. Gradient magnetic field being the most dominant parameter to manipulate MNPs in vitro and in vivo, it is possible to optimize this parameter using different types of magnet arrays as demonstrated in this study and other similar studies [Barnsley 2016].

Such numerical studies will help us to understand the principles behind time-varying magnet array induced cytotoxicity and facilitate the optimization of this process. From a numerical perspective, the size and magnetic susceptibility of the nanoparticles used (Polymag, nTmag - 100nm), the distance between the magnet array and the cell culture plate (1 mm), the rapid accumulation of MNPs by the use of an alternate pole magnet array, and the variation in magnetic force mediated by a time varying field gradient are behind the increased cytotoxicity of cancer cells.

## V. CONCLUSIONS

Therefore, from our field measurements, numerical calculations and our cytotoxicity experiments, surface functionalized, super paramagnetic iron oxide nanoparticle at low concentrations was externally manipulated, using a time varying field gradient technique (at very low frequencies, i.e., 1 to 4 Hz), to induce magneto-mechanical cell death in cancer cells. This is a key to the effective inducement of non-invasive cytotoxicity in cancer cells in physiological and clinical settings; further multidisciplinary research is required to make substantial progress in this area.

## ACKNOWLEDGMENT

MS worked for nanoTherics limited, but currently funded by the EPSRC and the Imperial College London. JD is a consultant for nanoTherics Limited. AM, SIJ and JL are academic researchers.

